# Remodeling of nucleosome by a DNA translocating bacterial restriction-modification enzyme

**DOI:** 10.1101/2025.01.15.633181

**Authors:** Sujata S. Gaiwala Sharma, Mahesh Kumar Chand, Ishtiyaq Ahmed, Nibrasul Haque, Kayarat Saikrishnan

## Abstract

A eukaryotic cell has specialized ATP-dependent chromatin remodelers, such as SWI/SNF, to unfurl DNA from nucleosome for functional processing. The ATPase that powers the movement of the chromatin remodeler on the DNA (translocation) is evolutionarily related to those powering the translocation of the functionally distinct bacterial restriction-modification (RM) enzymes. The collision of a SWI/SNF chromatin remodeler and a nucleosome, results in sliding/ejection of the constituent histone octamer, while two converging ATP-dependent RM enzymes catalyze DNA cleavage. Here, we investigate if an ATP-dependent Type ISP RM enzyme, an active and directional translocase, can remodel nucleosomes. Our results reveal that in presence of ATP, a Type ISP RM enzyme can displace the octamers from not just mononucleosomes but also two tandem nucleosomes. However, a Type III RM enzyme, which employs a homologous ATPase as a switch to facilitate bidirectional 1D diffusion along the DNA, fails to remodel the nucleosome. This implies that an actively translocating Type ISP RM enzyme generates sufficient force for chromatin remodeling, and may serve as artificial sequence-specific chromatin remodelers.

## Introduction

An oft-encountered feature of protein domains is their presence in functionally unrelated proteins across different kingdoms of life. One such domain is the ATPase belonging to the Superfamily 2 of helicases (1). These ATPases are often part of distinct proteins that process nucleic acids in diverse manner. The SF2 ATPases that interact with dsDNA are of three kinds – helicases that unwind double-stranded DNA (dsDNA), translocases that facilitate directional movement along dsDNA, or switches that facilitate bidirectional diffusion along the dsDNA. A striking example of the use of the ATPase for distinct purposes is i) to power endonucleolytic DNA cleavage by the bacterial ATP-dependent RM enzyme and ii) for nucleosome remodeling by the eukaryotic ATP-dependent chromatin remodelers. Here, we ask if the bacterial ATPase-containing RM enzyme, due to the presence of the conserved SF2 ATPase, can remodel nucleosome.

The SF2 ATPases in bacterial ATP-dependent RM enzymes power the nucleases that destroy invading foreign DNA. There are three types of ATP-dependent RM enzymes – Type I, Type ISP and Type III. A common feature of these enzymes is the requirement of at least two recognition sites on the DNA, which can be separated by as many as few thousand base pairs, for cleavage. In Type I/ISP RM enzymes, the SF2 ATPase function as translocases that bring the enzymes together in cis subsequent to binding to their respective recognition sites (2–4). The Type ISP enzymes (Fig. 1a), translocate at the speed of ∼300 bp/s (5) and it is their convergence that causes dsDNA break. In Type III RM enzymes (Fig. 1b), the SF2 ATPase acts as a switch that facilitates bidirectional 1D diffusion along the DNA, and cleavage results when an ATPase activated enzyme encounters another ATPase activated and site-bound enzyme (6,7).

**Figure 1.**
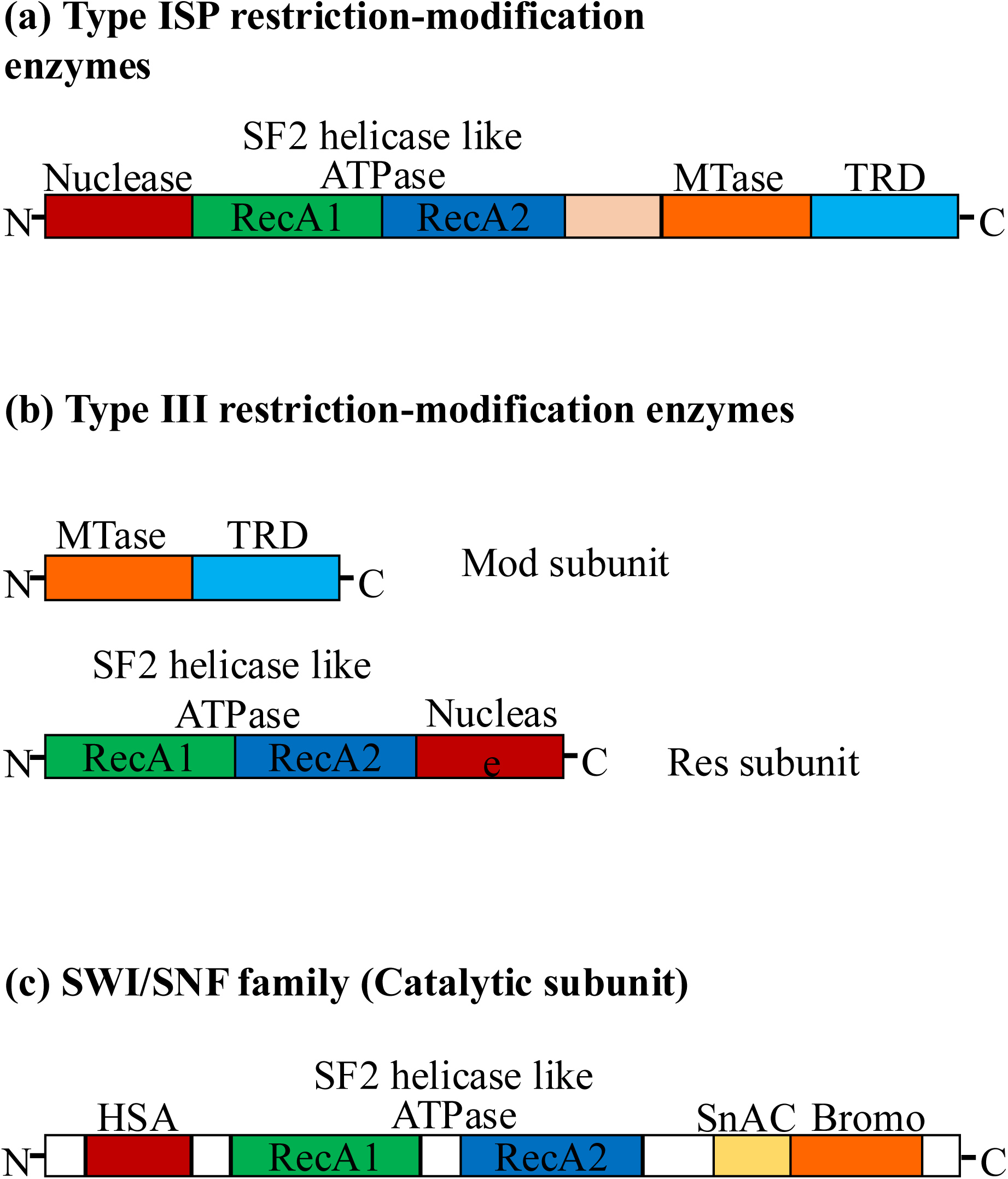
Domain architecture of a Type ISP RM enzyme, Type III RM enzyme and the catalytic subunit of SWI/SNF. A schematic of the domain arrangement in (a) Type ISP (b) Type III RM enzymes (Mtase - methyltransferase, TRD - Target Recognition Domain, Mod - modification subunit, Res - restriction subunit) and (c) SWI/SNF family of chromatin remodelers (HSA - Helicase-SANT-associated, SnAC - Snf2 ATP coupling).

In contrast to the RM enzymes, the SF2 ATPase in eukaryotic chromatin remodelers power nucleosome remodeling. In a eukaryotic cell, the dilemma associated in accommodating a 2-meter-long chromosome inside a 20-micron cell nucleus is overcome by packaging the chromosomal DNA into chromatin made of nucleosome - a nucleoprotein complex of chromosomal DNA wrapped around a histone octamer. However, the packaged DNA needs to be accessed time-to-time by the machinery for DNA replication, repair and transcription, which is facilitated by ATP-dependent chromatin remodelers.

There are different mechanisms of ATP-dependent nucleosome remodeling to free the DNA (8,9). The chromatin remodelers SWI/SNF and RSC complexes remodel chromatin by sliding or ejecting histone octamers (8–10). In these two complexes, the inbuilt SF2 ATPase (Fig. 1c) hydrolyzes ATP to power translocation of DNA by the remodeler attached to the octamer in the nucleosome (11,12). The translocating remodeler can travel at a speed of 25 bp/s and generate a force of 30 pN (13). The force thus generated disrupts the interaction between the octamer and DNA. It is proposed that the translocating remodeler bound to a nucleosome acts as a supercomplex that pulls the neighboring nucleosome dislodging the octamer on collision (13,14).

Though functionally distinct, the common feature between a Type I/ISP RM enzyme and a chromatin remodeler is the use of SF2 ATPase for high-speed translocation on DNA, which results in force generation. As the force generated by a remodeler can slide/eject the histone octamer, we wondered if a translocating RM enzyme could do the same. If capable, these enzymes could serve as a platform towards designing recognition sequence specific chromatin remodelers. The artificial remodeler would bind to a specific recognition sequence in the DNA and remodel the chromatin structure downstream, which we envisage can be used for targeted chromatin rearrangements *in vitro* and *in vivo*.

We report here our study on the ability of a Type ISP RM enzyme, LlaBIII and its nuclease-deletion mutant in remodeling both mono- and dinucleosome. We have also tested the nucleosome remodeling activity of EcoP1I, an enzyme belonging to the Type III RM enzyme family that diffuses on DNA. Our results indicate the importance of an active translocating ATPase motor of LlaBIII can remodel nucleosomes.

## Materials and methods

### Preparation of DNA substrates for nucleosome assembly

The DNA substrate used for *in vitro* nucleosome preparation was generated by PCR amplification. A 329 bp DNA was generated by amplification from a plasmid encoding the high affinity nucleosome occupancy region Widom (601) sequence (15) using primers 329F and 329R.The 329F primer incorporated LlaBIII and EcoP1I target sequence upstream of the 601. This fragment was used as template for incorporation of an adaptor sequence using primers Cy5oligo-498-AdaptorF (Cy5-F) and 329R. This produced a 320 bp DNA fragment that was used as a template for amplification of the 5’-Cy5 labelled DNA substrates. For terminally placed nucleosome DNA, 265 bp DNA was amplified using primers Cy5-F and 601R. For centrally placed nucleosomes, a 356 bp DNA was generated using primers Cy5-F and 329R. For unlabeled dinucleosome DNA, two fragments each containing a 601 region were generated using 329 bp fragment as template. The first fragment was generated using primers 329F and R1 overlap. This incorporated a NdeI site at the 3’ terminal of this DNA fragment. The second fragment was generated using primers F1 overlap and 329R. This incorporated a NdeI site at the 5’ terminal in this fragment. Both the fragments were purified using a PCR Cleanup Kit (Qiagen, Cat. No. 28106) and subjected to digestion by NdeI. The digested fragments were again purified using the PCR Cleanup Kit. Equimolar ratio of both the DNA fragments were ligated using the T4 DNA ligase (New England Biolabs, Cat. No. M0202S) according to the manufacturer’s specifications. Upon ligation, the fragments were resolved on a 1% agarose gel, stained with ethidium bromide, and visualized under a UV transilluminator. The band corresponding to the 498 bp ligated product was excised from the gel and purified using a gel extraction kit (Qiagen, Cat. No. 28704). The purified 498 bp DNA was utilized as a template for further PCR amplification using primers 329F and 329R, to generate more DNA. All the PCR products were purified using PCR Cleanup Kit (Qiagen, Cat. No. 28106) followed by purification from native polyacrylamide gel. The extracted DNA was again purified using the PCR clean up kit. The final DNA substrates were eluted in refolding buffer (2 M NaCl, 10 mM Tris pH 7.5, 1 mM EDTA, 10 mM β-mercaptoethanol) and concentration was estimated using NanoDrop™ 2000 (Thermo Scientific). The substrates were stored at -30°C until nucleosomes were reconstituted.

For preparation of the methylated DNA, the unmethylated 498 bp DNA was used as a template. Instead of the unmethylated cytosine, a dNTP mix with 5-methylcytosine was used in the PCR reaction. This led to amplification of a fully methylated 498 bp DNA, except for the ends where primers are incorporated, which were hemi-methylated. The methylated 498 bp DNA was also purified from native polyacrylamide gel as previously described and stored in -30°C until further requirement.

### Purification of human histones

The plasmids carrying the human histone encoding genes were acquired from Addgene (Catalog no.42634, 42630, 42633 and 42632). The histones were purified using the Rapid Histone Purification (RHP) protocol as published by (16) with minor changes wherever required (Supplementary Fig.1). Individual human histones were purified under denaturing conditions and ultimately stored as lyophilized powders at -30°C until required for octamer assembly.

### Assembly and purification of octamer

The lyophilized powders of individual histones were separately dissolved in unfolding buffer (7 M Guanidium Hydrochloride, 10 mM Tris pH 7.5 and 10 mM DTT) and protein concentration was estimated. The histones were mixed in equimolar ratio and dialyzed against refolding buffer (2 M NaCl, 10 mM Tris pH 7.5, 1 mM EDTA, 10 mM β-mercaptoethanol) at 4°C overnight. The buffer was changed once during the dialysis. The dialyzed protein was recovered from the dialysis bag and centrifuged at 4°C to remove any precipitates. The protein sample was then applied onto Superdex 200 10/300 column (GE Healthcare) pre-equilibrated with refolding buffer and fractions were analyzed by SDS-PAGE on an 18% polyacrylamide gel. Fractions corresponding to the octamer, molecular weight (109.542 KDa), elute at 12 ml elution volume. These fractions were analyzed by SDS-PAGE (Supplementary Fig. 2) and those that showed presence of bands corresponding to all four histones on polyacrylamide gel were pooled and concentrated using Vivaspin2 sample concentrator at 4000 rpm at 4 °C in Eppendorf centrifuge 5801 R. The protein concentration was estimated by Bradford’s assay and the concentrated protein was stored at -80°C.

### Purification of LlaBIII WT and LlaBIIIΔN

Purification of LlaBIII WT was performed as described by (2). LlaBIIIΔN cloning and purification has been previously described in PhD thesis by Dr. Mahesh Kumar Chand. Shortly, 1-165 amino acids that constitute the nuclease domain of LlaBIII were deleted from the full length *llaBiii* to obtain the construct expressing LlaBIIIΔN. The truncation of the nuclease domain was obtained by PCR amplification of the *llaBiiiΔn* from the full length *llaBiii* as template using primers LBΔN-pHIS-F and LB-PHIS-R. The ∼4.3kb *llaBiiiΔn* fragment was cloned into NdeI and BamHI sites in pHIS17 vector through restriction-ligation cloning method and the clone having the correct sequence of *llaBiiiΔn* was further processed for protein expression. The protein was expressed in *E. coli* BL21AI cells induced with 0.2% arabinose and purified following the same protocol as described for LlaBIII WT. All the proteins were stored at -80°C until further use.

### Purification of EcoP1I^E916A^

Purification of EcoP1I^E916A^ was performed as described previously (6).

### Reconstitution of nucleosome particles

The nucleosomes were assembled in vitro by following the protocol given by (17). Briefly, the histone octamer and 601 sequence containing substrate DNA were mixed in various protein:DNA ratios. The volume was made up to 10µL with refolding buffer which contains 2 M NaCl. The mixture was incubated on ice for 30 minutes. After 30 minutes, 10 µL of 10 mM Tris pH-7.5 was added and mixed gently by tapping and again incubated on ice for 1 hour. This process was repeated three more times, stepwise adding 5 µL, 5 µL and 70 µL of 10 mM Tris pH-7.5, thereby diluting the NaCl concentration in a stepwise manner from 2 M to 0.2 M. After the last incubation on ice for 1 hour the reconstituted nucleosome samples were loaded onto a 6% native polyacrylamide gel that was pre-run at 150 V for 20 minutes at 4°C in 1X TBE. The nucleosome samples were resolved for 90 minutes at 150 V at 4°C in 1X TBE. The gel was then stained with ethidium bromide solution prepared in 1xTBE and imaged on Typhoon Biomolecular Imagers (Amersham).

### Restriction Enzyme Accessibility Assay (REAA)

All the REAAs were performed at 37°C in 1x Cutsmart buffer (NEB). For 5’-Cy5labelled nucleosomes, 3.75 nM nucleosome and 20 nM LlaBIII were added in the reaction mixture and incubated for 5 minutes on ice following which HhaI (New England Biolabs, Cat. No. R0139S) was added to the mix. The reactions were started by addition of 4 mM ATP and incubated at 37°C for 2 hours. For unlabeled methylated and unmethylated dinucleosomes, 10-15 nM nucleosome and 100 nM enzyme (LlaBIII or EcoPI^E916A^) were mixed on ice. The mixture was incubated on ice for 5-10 minutes. 0.5 µL HhaI (New England Biolabs, Cat. No. R0139S) or for methylated substrate, MseI (New England Biolabs, Cat. No. R0525S) was added to the relevant reactions. The reactions were started with addition of 2 mM ATP, and the samples were incubated at 37 °C for 2 hours. ATP (2 mM) was supplemented halfway through the incubation. At the end of the incubation, the reactions were stopped with 2 µL of 0.5 M EDTA and 0.8 µL of 10% SDS. 2 µL of 2 mg/mL Proteinase K was added and incubated at 37°C for 30 minutes. The reaction mix was heated at 65°C for 20 minutes before addition of 4 µL ST buffer (100 mM Tris pH-7.5, 40% w/v Sucrose) in 5’-Cy5 labeled substrates or 3 ul of 6X Purple loading dye (New England Biolabs) in unlabeled substrates. The samples were resolved on a pre-run 6% native polyacrylamide gel run at 150 V for 50-55 minutes in 1x TBE at room temperature. Gels were imaged on Typhoon Biomolecular Imagers (Amersham) or Typhoon Trio+ Variable mode imager (Amersham). All the graphs were plotted and statistically analyzed using GraphPad Prism 10 software.

## Results

### LlaBIII can remodel mononucleosomes

The nucleosome remodeling potential of LlaBIII was tested using the **<u>r</u>**estriction **<u>e</u>**nzyme **<u>a</u>**ccessibility **<u>a</u>**ssay (REAA) (18,19) (Fig. 2 c and d). Pure histone octamers (Supplementary Fig.2) were assembled on 5’-Cy5 labelled DNA substrates containing a 147 bp long high-affinity octamer binding sequence 601 (15). Two DNA substrates were designed with distinct positions of the 601 sequence. Terminally positioned nucleosome (265 bp) contained the 601 sequence at the 3’ terminal (Fig. 2a) whereas centrally positioned nucleosome (356 bp) contained a 601 region in the center of DNA leaving a 91 bp overhang at the 3’ end of the 601 sequence (Fig. 2b). A LlaBIII binding site (5’TNAGCC3’) was incorporated 48bp upstream from the 601 sequence in both these DNA substrates (Fig. 2 a and b). There is another LlaBIII target site located close to the 5’ end of the DNA, although this site is too close to the DNA end (11bp) to allow LlaBIII translocation and can be ignored for the purpose of these assays. The quality of the assembled nucleosomes was checked using EMSA to ensure that free DNA was minimal and that the histone octamers were positioned as intended in both the nucleosome substrates (Supplementary Fig. 3 a and b). Each of the two 601 sequences had a recognition site for the nucleotide-independent restriction enzyme HhaI (GCGC). Each nucleosome (3.75nM) was individually incubated with LlaBIII (20 nM) for 5 minutes followed by addition of HhaI. Addition of ATP (4 mM) initiated the reaction. The nucleosome remodeling activity was monitored, in absence and presence of ATP, through cleavage of the DNA substrate by HhaI. Nucleosome displacement leading to exposure of HhaI target site and DNA cleavage was expected to yield two fragments in both of the nucleosome substrates. The 5’-Cy5 labelled DNA fragment in both the nucleosomes was expected to be 190 bp upon DNA cleavage (Fig. 3 a and d). The polyacrylamide gels were imaged using Cy5 specific wavelength filters, hence only the labelled fragment was visualized. Native PAGE analysis showed that in the absence of either LlaBIII or ATP, only the free DNA in the nucleosome preparation was cleaved whereas majority of the DNA that was nucleosome occupied remained unaffected by the presence of HhaI (Fig. 3 b-c and e-f, Lane 4 & 5). However, addition of HhaI, LlaBIII and ATP resulted in a drastic increase in the percentage of DNA cleaved in both reactions, with terminally placed and centrally placed nucleosomes (Fig. 3 b-c and e-f, Lane 6). Cleavage at HhaI sites indicated ATP dependent displacement of nucleosome by LlaBIII. Nucleosome displacement was also unaffected by the presence of an extra 91 bp overhang downstream of the 601 region in the centrally positioned nucleosome. Irrespective of the presence or absence of downstream DNA overhang LlaBIII could remodel the mononucleosomes.

**Figure 2.**
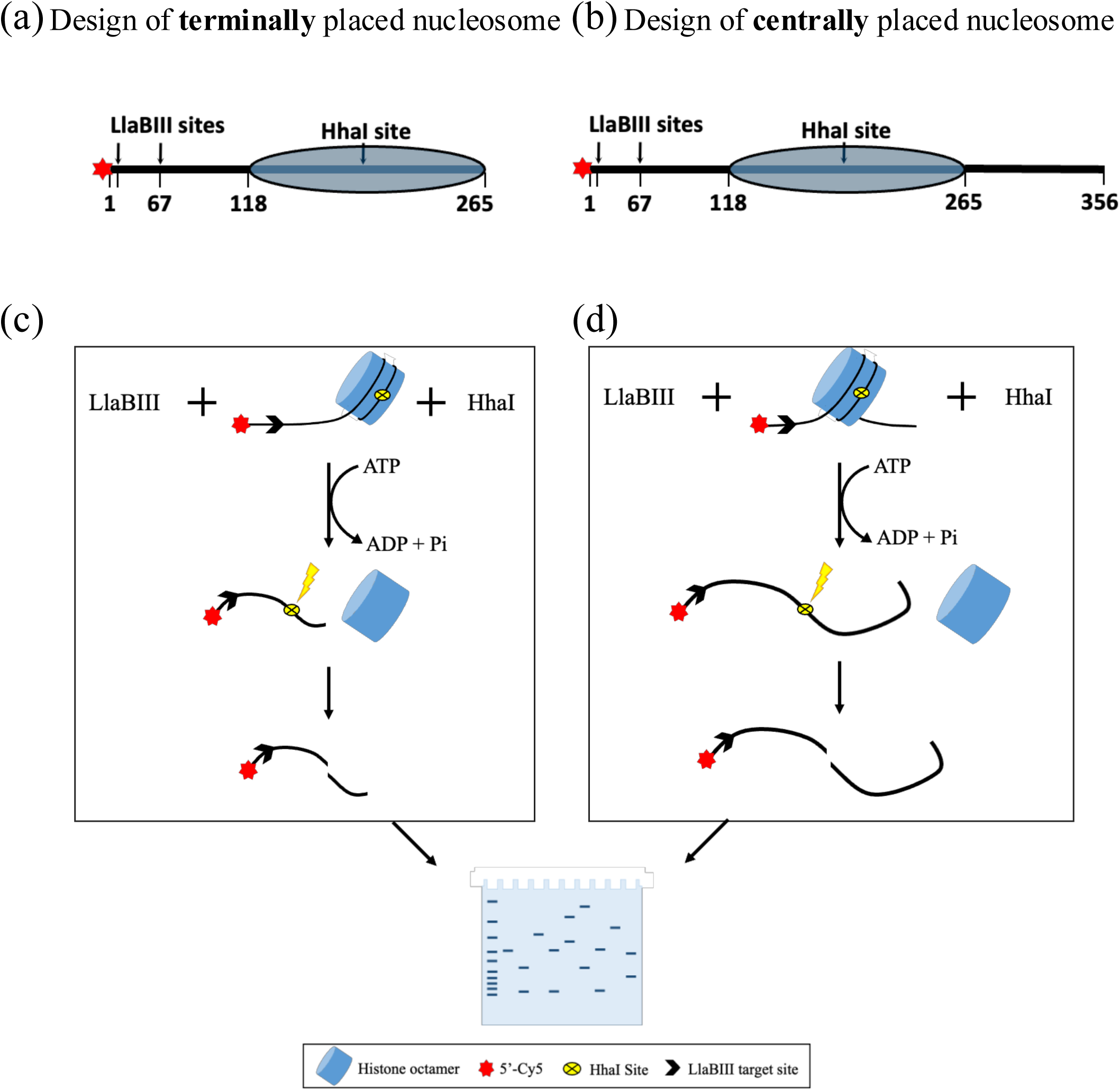
A graphic representation of the restriction enzyme accessibility assay (REAA). (a-b) Graphical representation of terminally placed and centrally placed mononucleosomes. Blue ovals represent the octamer binding region, 601 and black lines depict DNA. The numbers specify base pair positions of the indicated features. (c-d) Graphical illustration of REAA with terminally placed and centrally placed mononucleosomes.

**Figure 3.**
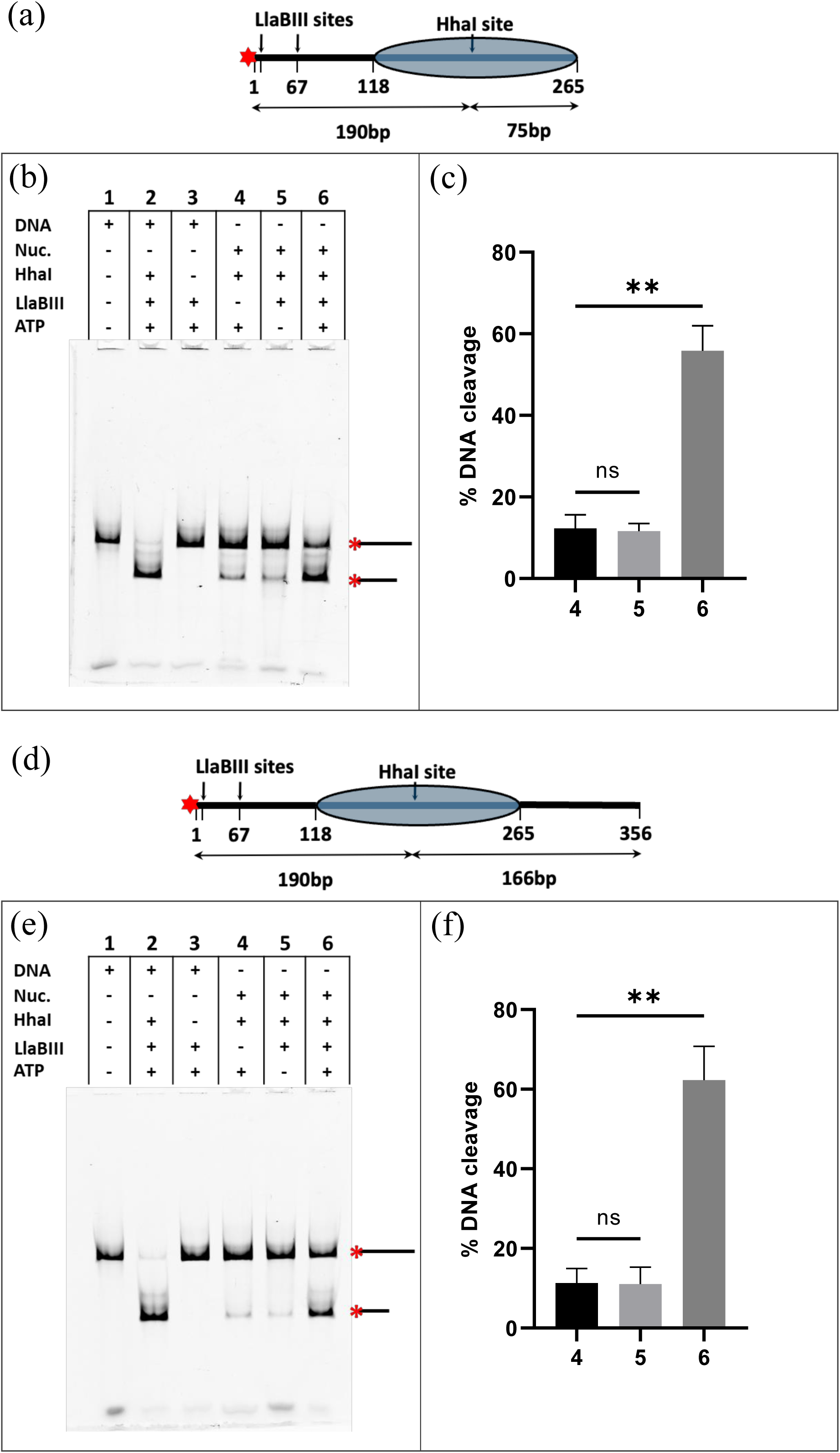
Nucleosome remodeling by LlaBIII. (a and d) Graphical representations of terminally placed and centrally placed mononucleosomes. Blue ovals represent the octamer binding region, 601 and black lines depict DNA. The numbers specify base pair positions of the indicated features. Arrows indicate the expected DNA fragments upon HhaI cleavage. Red star indicates 5’-Cy5 label. (b-c) REAA with terminally placed nucleosome and LlaBIII. (e-f) REAA with centrally placed nucleosome. In both the nucleosomes, HhaI site is inaccessible in presence of octamer and hence DNA is protected as seen in lanes 4. LlaBIII in absence of ATP does not affect the nucleosome as seen in lanes 5. Whereas, in presence of ATP, LlaBIII displaces the octamer and makes the DNA accessible to cleavage by HhaI as observed from lanes 6. The bar plots show the percentage nucleosomal DNA cleaved by HhaI in absence of LlaBIII (4), in presence of LlaBIII (5) and in presence of LlaBIII and ATP (6), N=3. Statistical significance was assessed using the unpaired t-test (*P <0.05, **P <0.01, ***P <0.001), N≥3.

### LlaBIII can displace both octamers from a dinucleosome substrate

Further, we introduced another 601 sequence on the nucleosome DNA separated from the first 601 sequence by a 22bp linker DNA (Fig. 4a). This allowed assembly of two nucleosomes on the DNA upon reconstitution. A LlaBIII recognition sequence (TGAGCC) was present 54 bp upstream of the first 601 sequence (Fig. 4a). The quality of the assembled dinucleosome was checked using EMSA and REAA to ensure that free DNA was minimal and that the histone octamers were positioned properly (Supplementary Fig. 4). Each of the two 601 sequences had a recognition site for the nucleotide-independent restriction enzyme HhaI (GCGC). The dinucleosome (10-15 nM) was incubated with LlaBIII (100 nM) in absence and presence of ATP (2mM). The remodeling activity was monitored through cleavage of the 498 bp unlabeled substrate DNA by HhaI. Exposure of and cleavage at either of the two HhaI sites was expected to yield ∼332 bp and ∼166 bp fragments, while cleavage at both the sites was expected to result in three fragments ∼163 bp, ∼169 bp and ∼166 bp in length (Fig. 4a).

**Figure 4.**
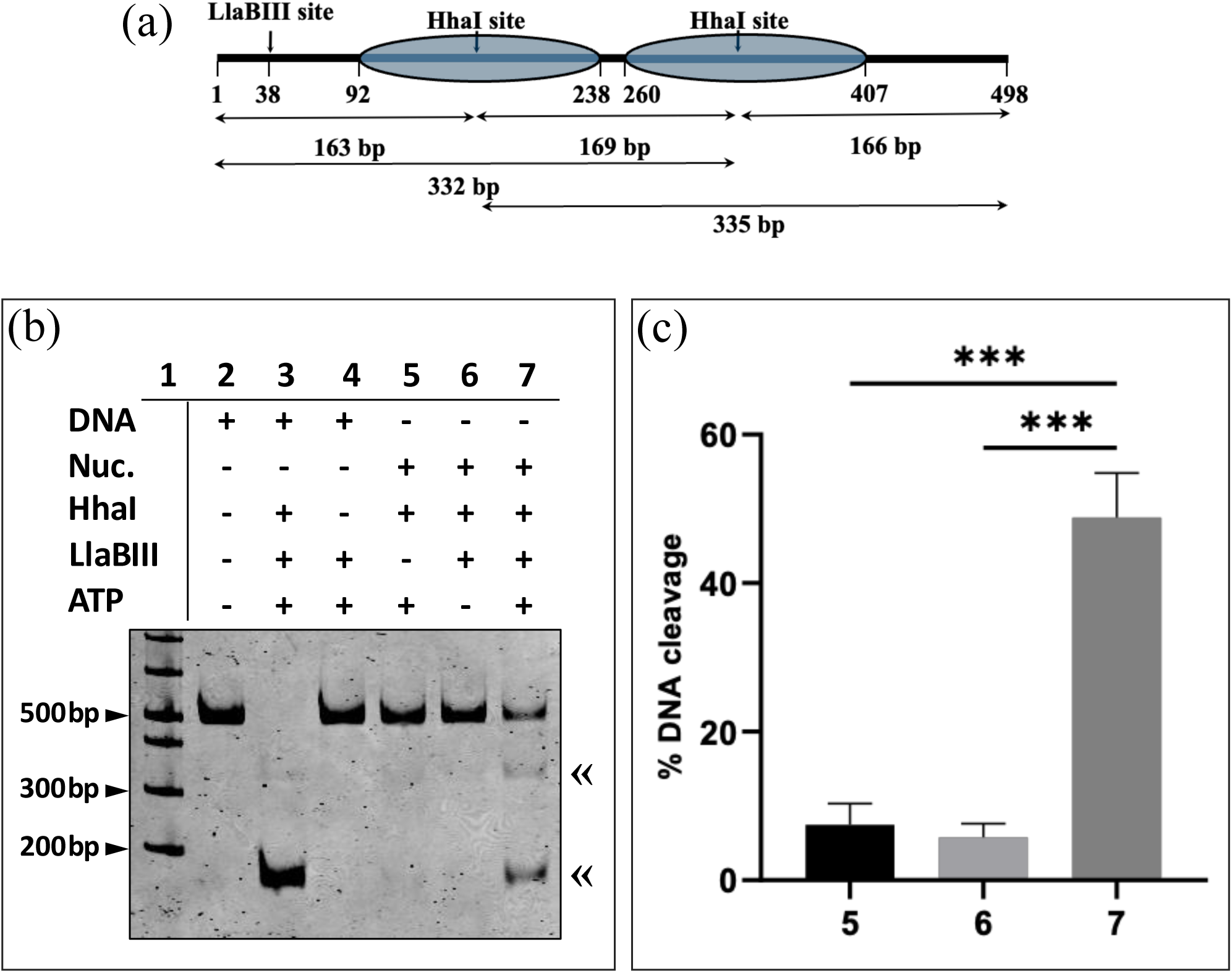
REAA with dinucleosome. (a) Schematic showing the positions of the HhaI target sites and the fragments generated upon DNA cleavage by HhaI. Gray ovals represent nucleosome occupancy region. (b) Representative native PAGE image accompanied by (c)-bar plots with percentage of DNA cleavage plotted on the Y-axis with corresponding lane numbers from the gel on X-axis. The arrows («) mark the positions of the expected DNA cleavage products on the polyacrylamide gel. Error bars represent standard error mean for four separate trials. Statistical significance was assessed using the unpaired t-test (*P <0.05, **P <0.01, ***P <0.001), N≥4.

When resolved on a native PAGE, we observed that either in the absence of LlaBIII or ATP, the integrity of the DNA remained unaffected by the presence of HhaI (Fig. 4 b-c, Lane 5 & 6). However, addition of LlaBIII, ATP and HhaI resulted in a prominent ∼166 bp cleaved DNA fragment and a less prominent fragment that migrated just above 300 bp (Fig. 4 b-c, Lane 7). The ∼166 bp band was consistent with the DNA fragments obtained when HhaI cleaved at both its sites. As the size of the three fragments differed by only a few base pairs, they failed to be resolved into individual bands. The band near 300 bp was interpreted as the ∼332 bp fragment produced when HhaI cleaved only at one site. This result clearly indicated that in the presence of LlaBIII and ATP the two HhaI sites were exposed for cleavage. The occurrence of the ∼300 bp DNA fragment was suggestive of step-wise remodeling.

### Deletion of nuclease domain does not affect remodeling activity of LlaBIII

We also tested if a deletion of the N-terminal nuclease domain of LlaBIII (LlaBIIIΔN) affected its ability to displace the histone particles. We had earlier shown that a similar nuclease-deleted construct of LlaGI, a close homologue of LlaBIII sharing ∼80% sequence identity, is an active ATPase and translocase (3). We performed remodeling assays with LlaBIIIΔN and found that it was capable of nucleosome remodeling with similar efficiency as the wild-type enzyme (Fig. 5 b-c, Lane 6 &7).

**Figure 5.**
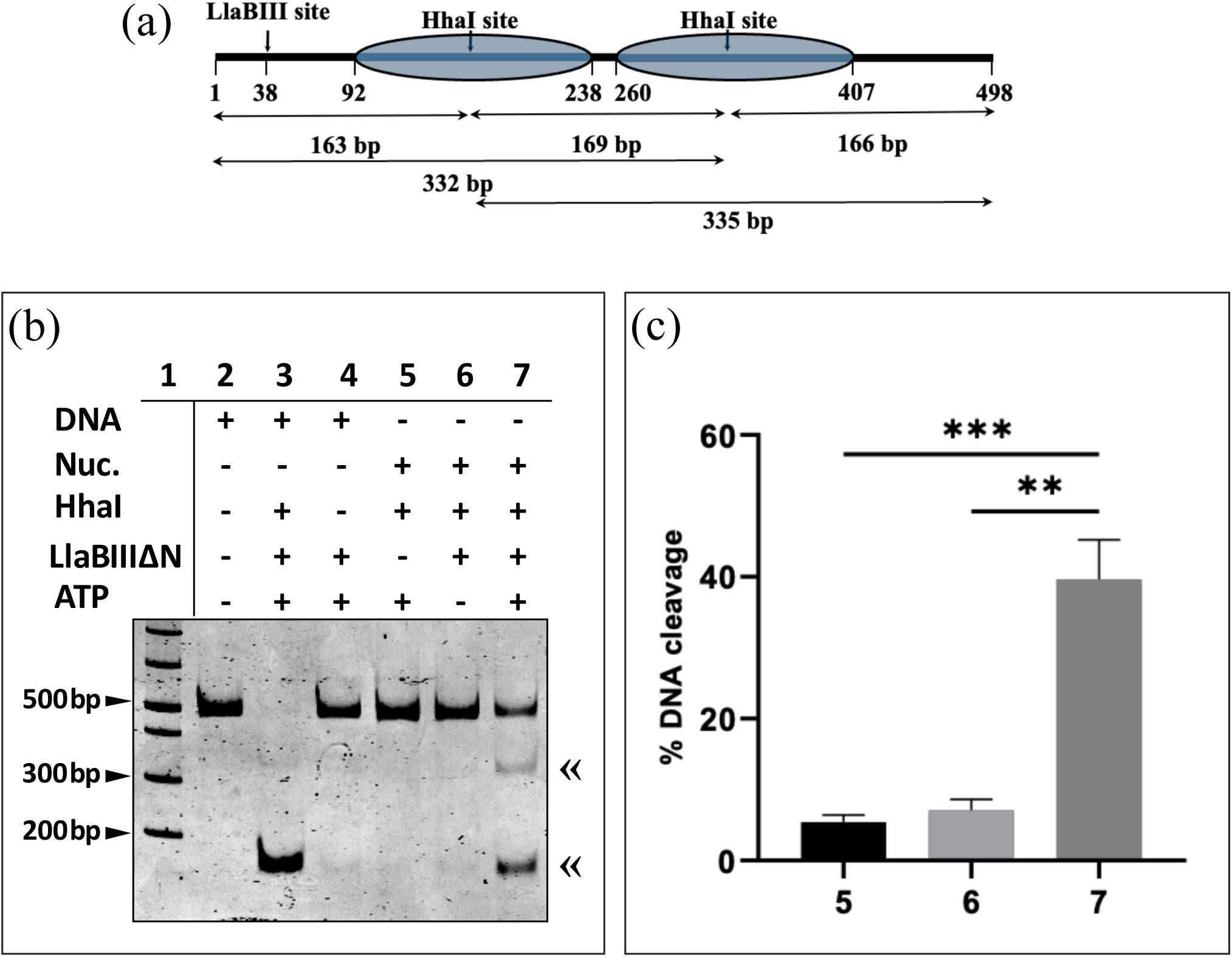
REAA with dinucleosome and LlaBIIIΔN. (a) Schematic showing the positions of the HhaI target sites and the fragments generated upon DNA cleavage by HhaI. Blue ovals represent nucleosome occupancy region. (b) Representative native PAGE image accompanied by (c)-bar plots with percentage of DNA cleavage plotted on the Y-axis with corresponding lane numbers from the gel on X-axis. The arrows («) mark the positions of the expected DNA cleavage products on the polyacrylamide gel. Error bars represent standard error mean for four separate trials. Statistical significance was assessed using the unpaired t-test (*P <0.05, **P <0.01, ***P <0.001), N≥4.

### A diffusing SF2 ATPase does not remodel the dinucleosome

Having established that the Type ISP enzyme can dislodge histone octamers, we next asked if the SF2 ATPase in a Type III RM enzyme could do the same. The SF2 ATPase in Type III RM enzymes is not a translocase, and instead functions as a switch facilitating the 1D diffusion of the enzyme along the DNA and activating the nuclease (6,7). We performed REAA using the dinucleosome substrate, which had a recognition sequence of the Type III RM enzyme EcoP1I, AGACC, adjacent to the LlaBIII recognition sequence (Fig. 6a). A nuclease-dead but ATPase-active mutant of EcoPI (EcoP1I^E916A^) was used for these experiments to prevent the enzyme from catalyzing the non-canonical single-site DNA cleavage (6). The REAA experiment clearly showed that, unlike LlalBIII, EcoP1I^E916A^ did not remodel the dinucleosome and expose the HhaI sites for cleavage even in the presence of ATP (Fig 6 b-c, Lane 5-7).

**Figure 6.**
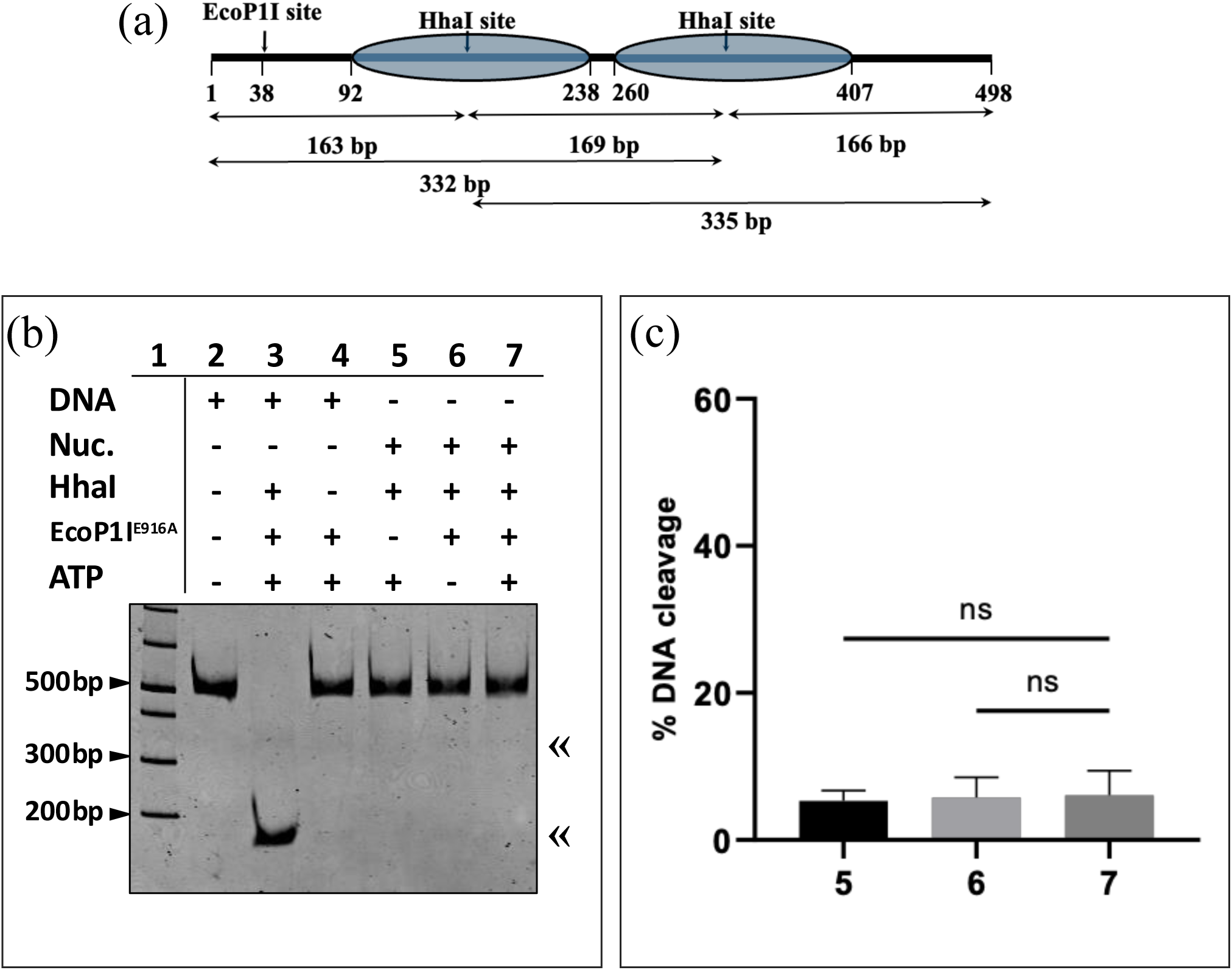
REAA with dinucleosome and EcoP1I^E916A^. (a) Schematic showing the position of EcoP1I and HhaI sites and sizes of the DNA fragments that would be produced upon cleavage by HhaI. Blue ovals depict the nucleosome occupancy region. (b) Representative native PAGE image accompanied by (c)-bar plots with percentage of DNA cleavage plotted on the Y-axis with corresponding lane numbers from the gel on X-axis. The arrows (**«**) mark the positions of the expected DNA cleavage products on the polyacrylamide gel. Error bars represent standard deviation for two separate trials. Statistical significance was assessed using the unpaired t-test (ns P > 0.05), N=2.

### Cytosine methylation does not hinder nucleosome remodeling by LlaBIII

In a cell, the DNA is almost always methylated, and the nucleosomes are always in association with the DNA containing methylated cytosine. In literature, there are contradictory reports about how the presence of methylated cytosines in the DNA affects the nucleosome structure in terms of the compaction, stability, and occupancy of nucleosome (20–26). Nevertheless, we decided to check whether remodeling of nucleosomes prepared from methylated DNA display any differences in the nucleosome remodeling. For this we prepared dinucleosomes with cytosine methylated DNA and performed REA assay.

REAA with methylated nucleosome utilized the two MseI restriction sites situated approximately in the center of the nucleosome occupying region (601) (Fig. 7a). Cleavage by MseI is unaffected by cytosine DNA methylation, hence could be utilized in the REA assay instead of HhaI. We found that both LlaBIII and LlaBIIIΔN could displace nucleosomes from the methylated DNA in presence of ATP (Fig. 7 b-e). The extent of nucleosome displacement at the end of two hours was similar to that observed for unmethylated nucleosome. This suggested that the interactions between the DNA and octamer might remain unaltered or increase upon cytosine methylation. We also tested if EcoP1I^E916A^ could remodel the methylated dinucleosomes. We observed that there was no indication of nucleosome displacement from the methylated DNA by EcoP1I^E916A^ (Fig.7 f-g). This indicated that the interaction between a methylated DNA and histone octamer was not significantly weakened to allow a DNA diffusing enzyme like EcoP1I^E916A^ to displace the octamer from its position on the DNA.

**Figure 7.**
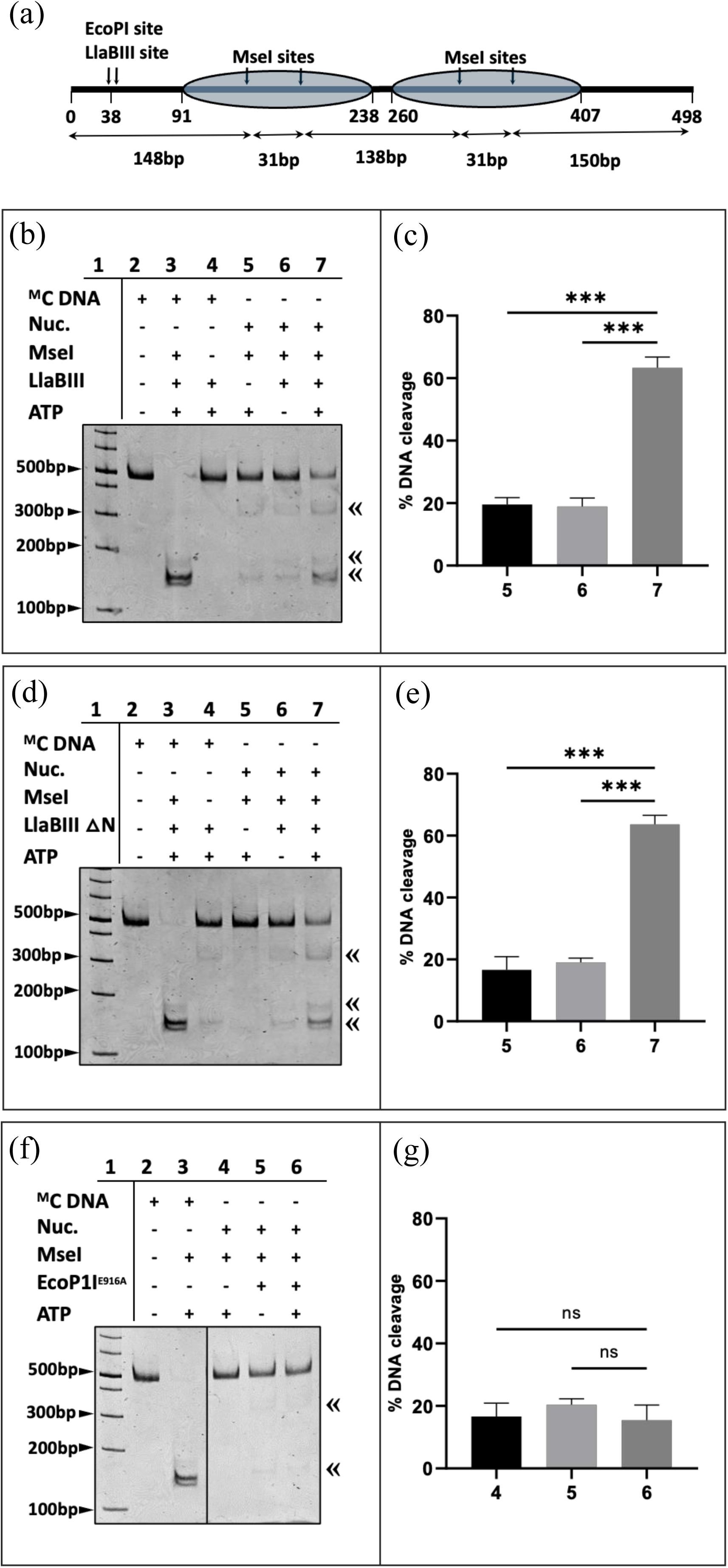
REAA with methylated dinucleosome. (a) Schematic showing the design of the dinucleosome assembled on methylated DNA. Blue ovals represent the nucleosome occupying region (601). Two MseI sites in each 601 regions are marked and arrows depict the DNA fragments generated upon cleavage by MseI. Representative native PAGE images showing REAA with (b) LlaBIII (d) EcoP1I^E916A^ (f) LlaBIII^R564A^ (h) LlaBIIIΔN. The arrows (**«**) mark the positions of the expected DNA cleavage products on the polyacrylamide gel. The images are accompanied by bar plots with percentage of DNA cleavage plotted on the Y-axis with corresponding lane numbers from the gel image on X-axis. Statistical significance was assessed using the unpaired t-test (*P <0.05, **P <0.01, ***P <0.001), N≥3.

## Discussion

The Superfamily 2 (SF2) of helicases includes both the bacterial defense systems, such as the Type ISP RM enzymes, and the eukaryotic chromatin remodelers, such as the Snf2 family (1). Every member of these two families of proteins possesses the same conserved ATPase motor domain containing two RecA-like fold subdomains. Although these enzyme complexes perform strikingly different functions, the motor at the core performs the same function, that is, translocation on double stranded DNA. To achieve the varied DNA translocation related cellular functions, accessory domains or subunits are present along with the core motor. Here we show that LlaBIII, a prototype of the Type ISP family, can perform the basic functions of the chromatin remodelers like the Swi2/Snf2, by virtue of the conserved and modular motor domain. Like the chromatin remodelers, it can displace nucleosome from its position on a DNA. To demonstrate that the Type ISP RM enzymes can remodel nucleosome, we tested the nucleosome remodeling activity of LlaBIII using the *in vitro* assay – REAA. We have demonstrated that the Type ISP RM enzyme LlaBIII in presence of ATP can remodel mono- and dinucleosome to expose the masked HhaI restriction sites. Moreover, a minimal construct of the nuclease-deleted LlaBIIIΔN that had the SF2 ATPase and the modules required for sequence recognition, i.e. the methyltransferase and target recognition domain (2,3), was also proficient in remodeling. In contrast, the Type III RM enzyme (EcoP1I), in which the SF2 ATPase is a switch (activating 1D diffusion along the DNA) rather than an active translocase, failed to remodel the dinucleosome. Additionally, a common DNA modification present on nucleosomes in live cells, that is, presence of cytosine methylation on the nucleosome DNA does not inhibit or enhance the nucleosome remodeling activities of these enzymes. LlaBIII wild type and the nuclease truncated construct displace nucleosome as proficiently even with methylated nucleosomes whereas EcoP1I fails to affect nucleosome occupancy on this DNA as well. This is a critical characteristic of LlaBIII that allows for its potential development into an *in vivo* target dependent chromatin remodeler.

We interpreted our data to imply that the active translocase produced enough force to displace the histone octamers upon collision. Though a diffusing Type III RM enzyme may collide with the nucleosome, the force exerted does not appear to be sufficient for remodeling. These assays together demonstrated that the remodeling of the nucleosome required an SF2 ATPase, which was an active translocase.

We envisage that nuclease- and methylation-dead derivatives of these Type ISP RM enzymes can be useful tools in sequence-dependent remodeling of chromatin in vitro and in vivo. Silencing of the nuclease and methyltransferase activities would prevent unwanted nicking/cleavage or methylation of the substrate DNA, which could hinder site-specific remodeling. Furthermore, the large number of Type ISP RM enzymes with different recognition sequences occurring in nature could serve as a library of remodelers that are proficient in targeted chromatin remodeling.

