## Supplementary Figures for "Remodeling of nucleosome by a DNA translocating bacterial restriction-modification enzyme"

Supplementary figure 1

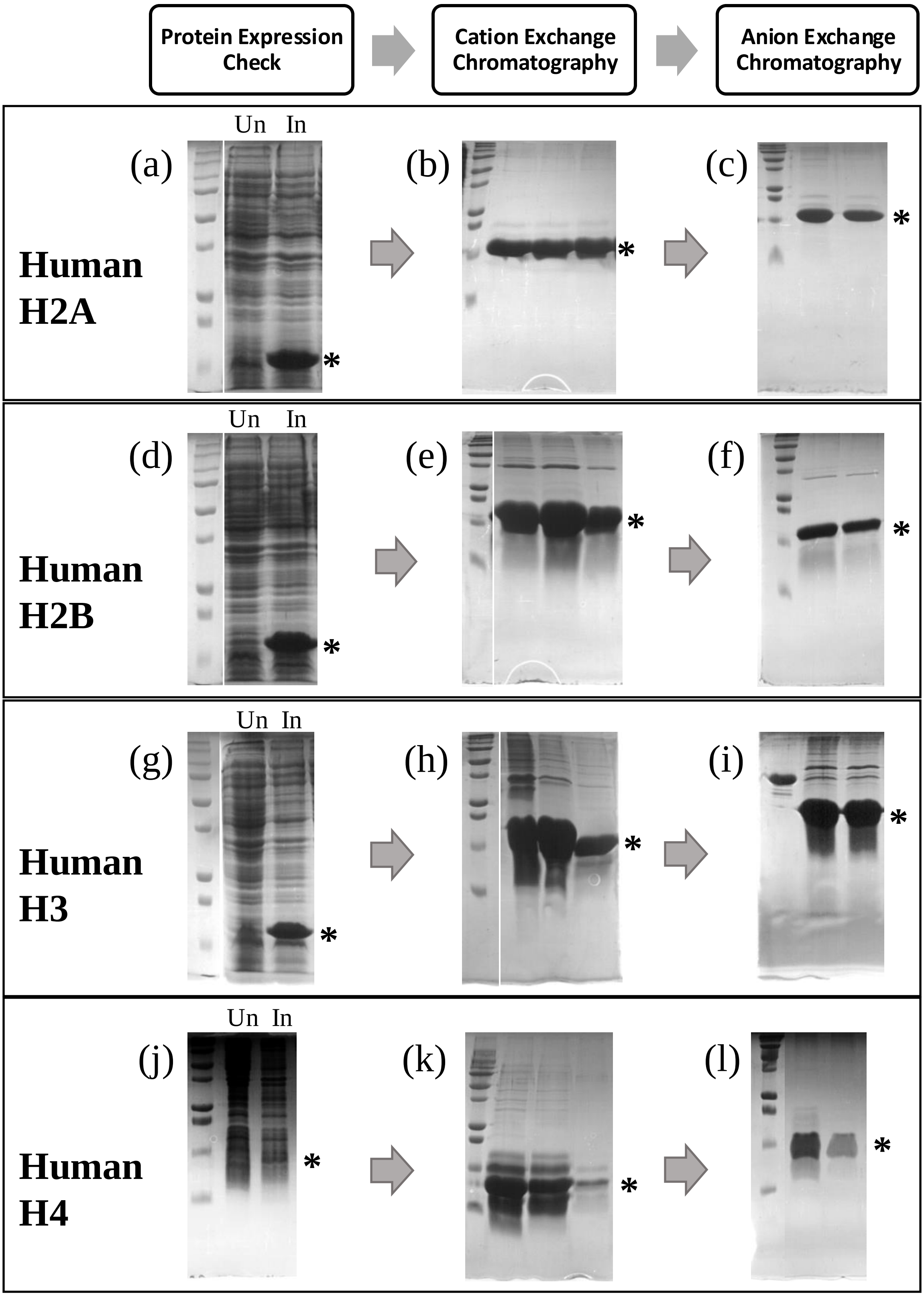


**Supplementary Figure 1**. Overexpression and purification of human histones H2A, H2B, H3 and H4 using the RHP protocol. (a)(d)(g) and (j) – SDS-PAGE gels showing overexpression of the respective histone proteins in *E. coli* BL21DE3. Asterisk (*) marks the position of the respective protein bands in the gels. (b)(e)(h) and (k) – SDS-PAGE gels showing fractions from cation exchange chromatography (HiTrap SP HP, GE Healthcare) containing the respective histone proteins. (c)(f)(i) and (l) - SDS-PAGE gels showing fractions from anion exchange chromatography (HiTrap Q HP, GE Healthcare) containing the respective histone proteins.

Supplementary figure 2

(a) Size exclusion chromatography profile

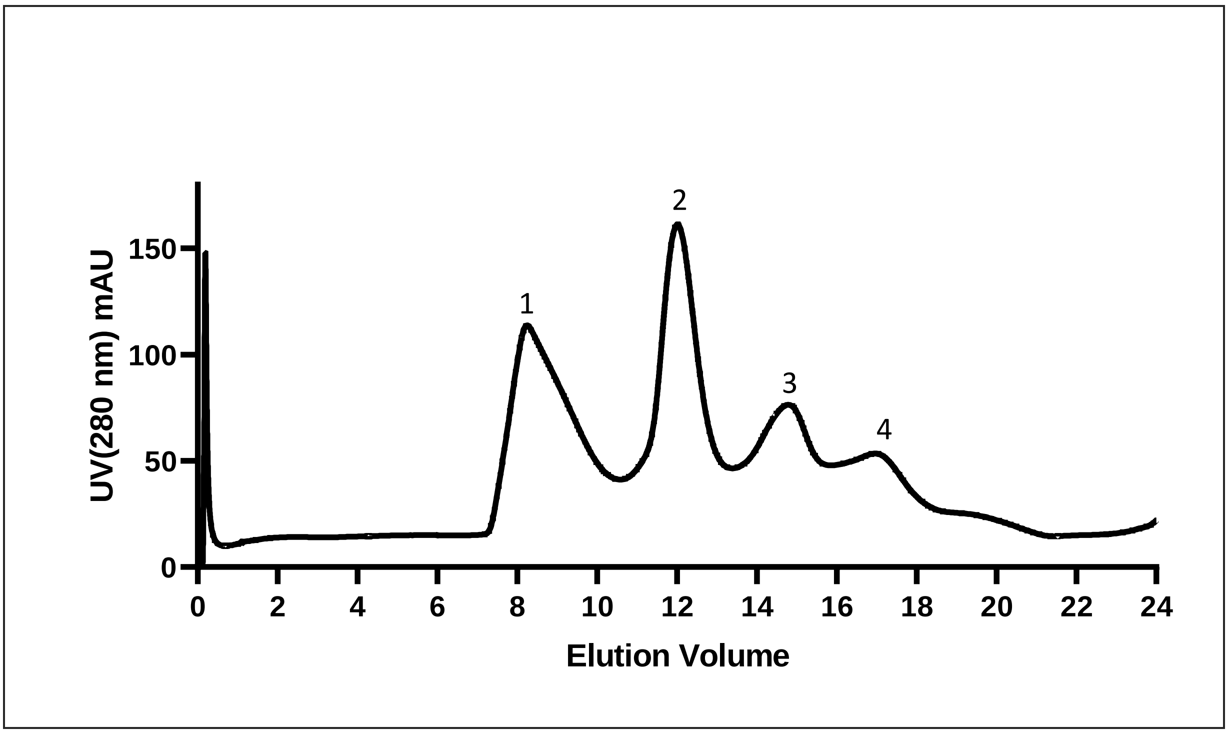


(b) SDS-PAGE-Fractions obtained from SEC

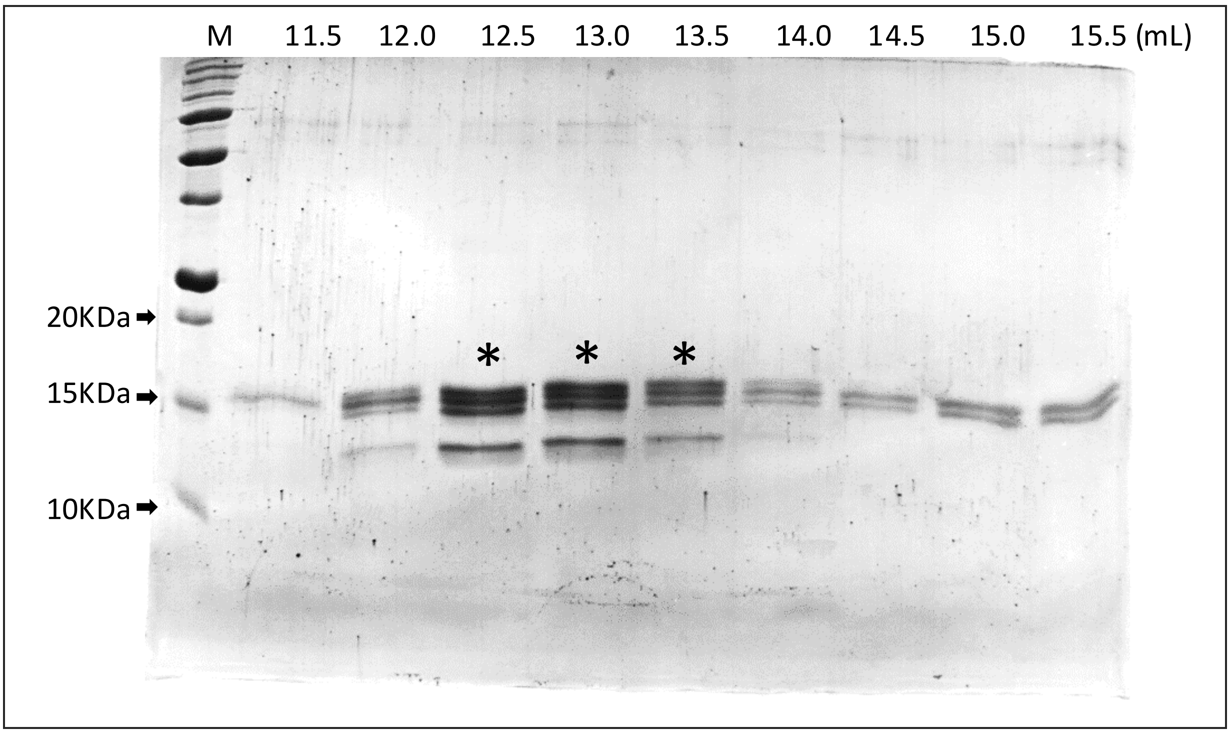


**H3** - 15.39 kD **H2A -** 14.09 kD **H2B -** 13.92 kD **H4 -** 11.37 kD

**Supplementary Figure 2**. Purification of assembled octamer. (a) Elution profile of refolded octamer resolved by size exclusion chromatography (SEC) (Superdex 200 10/300 column, GE Healthcare). 1 – contaminants (high molecular weight) (8.23 mL) 2 – refolded octamer (12.01 mL) 3 – H2A-H2B dimer (14.8 mL) 4 – contaminants (low molecular weight) (17.3 mL). (b) Fractions from SEC resolved by SDS-PAGE. Bands corresponding to each of the four histones are visible in the fractions corresponding to molecular weight of refolded octamer. Asterisk (*) marks the fractions pooled and concentrated for nucleosome reconstitution. N≥3

Supplementary figure 3

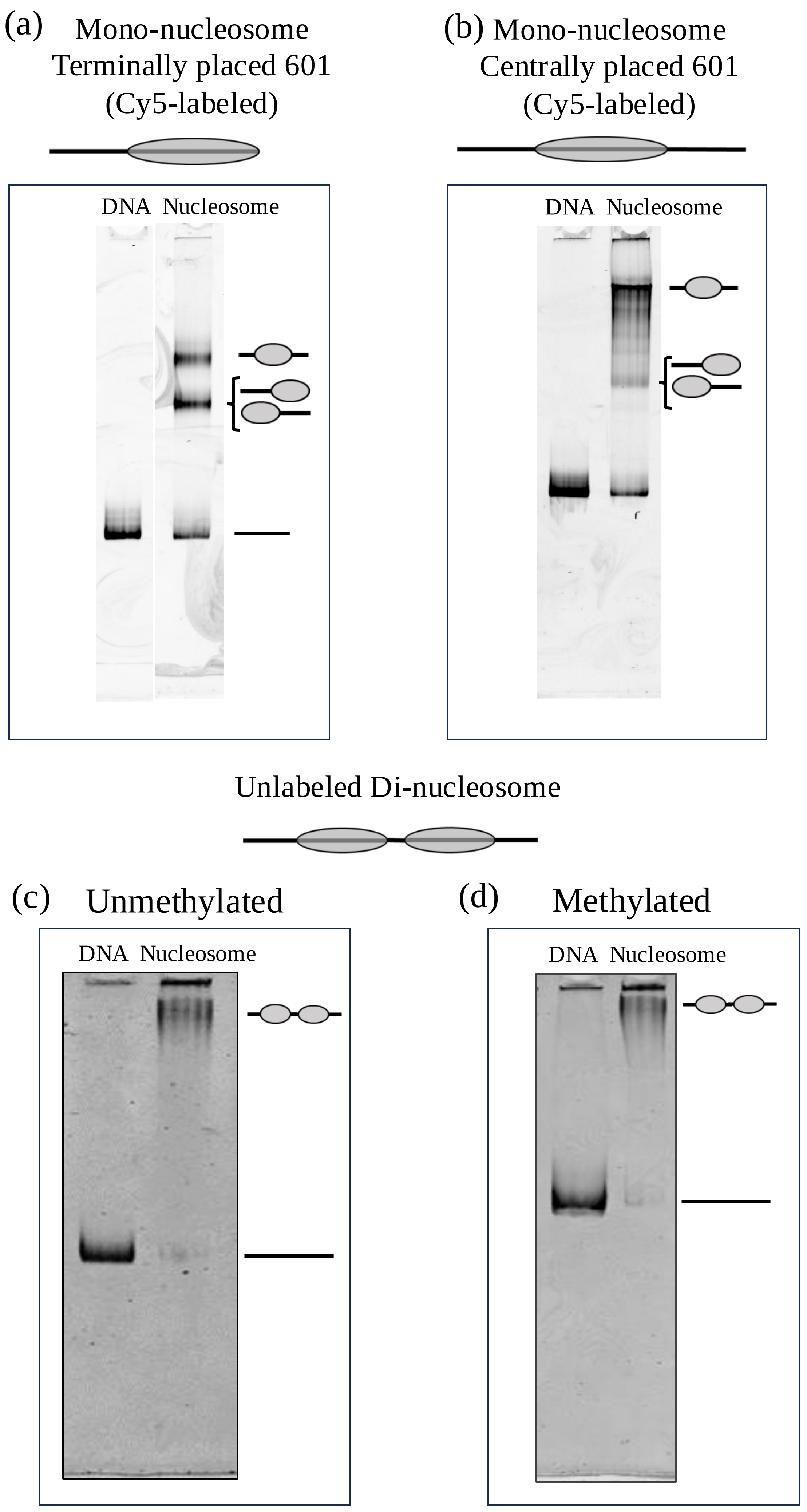


**Supplementary Figure 3**. Nucleosome positions observed on native polyacrylamide gel. (c) Mononucleosome assembled on Cy5 labelled 356 bp DNA with 601 sequence positioned centrally. (d) Dinucleosome assembled on Cy5 labelled 525 bp DNA carrying two 601 sequences.

### Supplementary figure 4

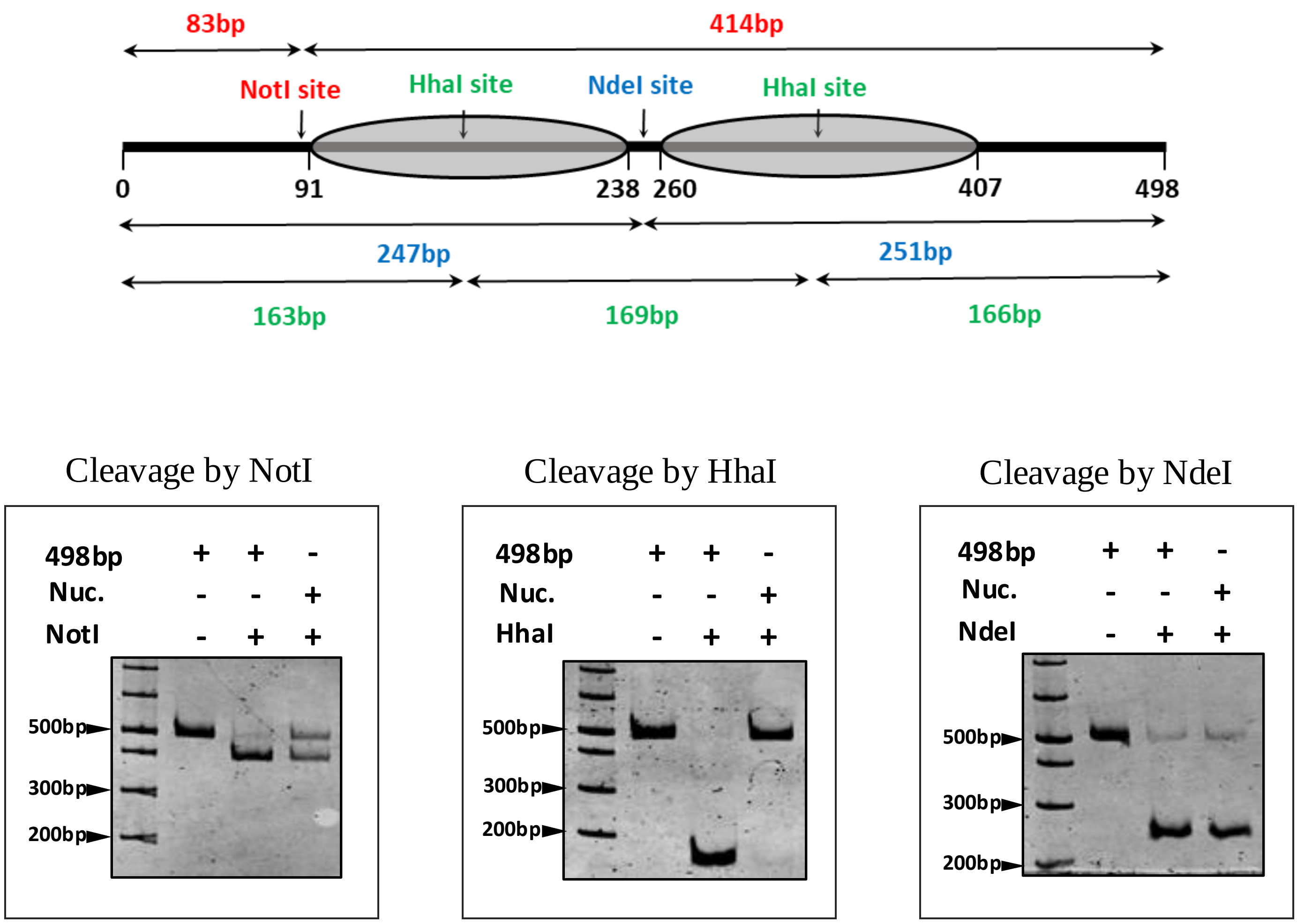


**Supplementary Figure 4**. Nucleosome positioning check. The nucleosome occupancy on the substrate DNA was confirmed by digestion with the specified restriction enzymes and the cleavage fragments were analyzed on native PAGE. (a) Schematic representation of the dinucleosome substrate depicting the position of target sites for NotI (Red), HhaI (Green) and NdeI (Blue), and the fragments generated upon cleavage by these enzymes individually. Grey ovals represent the nucleosome occupancy. Representative native PAGE images showing the DNA cleavage pattern for free DNA and dinuclesosome upon cleavage by NotI (b), HhaI (c) and NdeI (d). N≥2

### Supplementary table 1

#### DNA oligos used in the study

| Name | Sequence (5'-3') |
|---|---|
| 329F | CGAGGGGTAGGCCGCATCCTAATGACCTAATCAGACCTGAGCCTTCACACCG |
| 329R | CGAATTCGAGCTCGGTACCC |
| Cy5oligo-498-AdaptorF | CCCGTAGTGCTGTGCTGATAATCAGACCTGAGCCTTC |
| Cy5-F | TGGACACGACTTGAGCCCTCCTCACTTTCTTCCTCACCCGTAGTGCTGTGCTGA |
| 601R | ACAGGATGTATATATCTGACACG |
| F1 overlap | GCATCTGCATATGTATCCCGCCCTGGAGAATCCCGG |
| R1 overlap | GGCGGGATACATATGCAGATGCACAGGATGTATATATC |
| LBΔN-pHIS-F | CTTTAAGAAGGAGATATACATATGCGTCCAGAAAATGTTGTTG |
| LB-PHIS-R | CGCAGCAGCGGTTTCTTTACCAGACTCGAGTTATAGTCCCTGTACTACTCTTG |
